# Seroprevalence and risk factors associated with brucellosis in goats in Nyagatare district, Rwanda

**DOI:** 10.1101/2023.05.01.538860

**Authors:** Jean Paul Habimana, Jean Bosco Ntivuguruzwa, Aime lambert Uwimana, Aurore Ugirabe, Eric Gasana, Henriette van Heerden

## Abstract

Caprine brucellosis, normally caused by *Brucella melitensis* in goats, is endemic in Rwanda. However, reliable data on caprine brucellosis in Rwanda is insufficient and data on the prevalence and risk factors linked with goats’ brucellosis in the district of Nyagatare is unknown. A cross-sectional study was conducted among herds of goats in six sectors of Nyagatare district (*n* =102), to characterise factors at herd level associated with brucellosis seroprevalence in goats. Serum from goats was screened using both the indirect enzyme-linked immunosorbent assay (iELISA) and the Rose Bengal test (RBT). A tested systematic questionnaire was used to obtain data about major risk factors for brucellosis. Brucellosis seroprevalence was 6.8% and 10.7% on RBT and iELISA respectively. The overall seroprevalence was 6.8% on animal level and 16.6% on the herd level in series with RBT and iELISA. Mixing a herd of cattle and goats and history of abortions were the risk factors identified to be considerably linked with *Brucella* seropositive herd (*p* < 0.05). This study confirmed that brucellosis is endemic in the area, and a one-health strategy for controlling and preventing brucellosis in the Nyagatare district is strongly recommended. The study recommends an awareness campaign to educate all livestock farmers on brucellosis, further studies are recommended to characterize the *Brucella* spp. in small ruminants in Rwanda and recommend appropriate control measures.

## Introduction

Brucellosis is among the most widespread zoonotic infections in the world caused by *Brucella* species. *Brucella* bacterium is a rod-shaped Gram-negative (coccobacilli), non-motile (1, 2). Brucellosis is a herd disease affecting cattle that is normally caused by *B*.*abortus*, porcine brucellosis is caused by *B. suis*, and *B. melitensis*, which causes *Brucella* infection in goats and sheep, and it is very pathogenic to humans (2, 3). Brucellosis causes sterility and reproductive failure in livestock, and it often results in abortion and complete or delayed infertility in susceptible hosts (2, 4, 5). Infected animals disseminate *Brucella* agents in the colostrum and milk, it is also spread from uterus discharges ensuing abortion and subsequent parturition (6-9). The spread of brucellosis within and between herds is enhanced by different risk factors, including the introduction of new asymptomatically infected and non-quarantined animals (10). It is also amplified by the history of abortions within a herd (11-13). The mixing of different herds of animals during grazing or sharing watering places was also characterised as a risk factor (14, 15).

Many serological tests are used in the screening of *Brucella* infection in livestock. However, a single serology test is not suitable for every animal species and all circumstances. The positive results of the screening test ought to be confirmed with a confirmation serological test (2). Serological tests are not fully specific as they cross-react with other bacterial diseases, especially *Yersinia enterocolitica* O: 9 (16). The Rose Bengal Test (RBT) is a quick and easy agglutination test that uses an antigen that has been stained with rose bengal and buffered to a low pH of 3.65 0.05. (17). One of the significant brucellosis confirmation serological tests suggested for small ruminants is indirect or competitive enzyme-linked immunosorbent assays (iELISA & cELISA) using the smooth lipopolysaccharide (sLPS) antigen. (18-21).

Rwanda has an estimated 2.1 million agricultural households and livestock ownership is distributed as follows; 61.0% of households own cattle and 53.6 % with goats (22). Rwanda had an estimated number of 2,283,445 goats, 1,856,490 cattle, 703,145 pigs, and 499,316 sheep (22). Goat population decreased by 6.5%, which is attributed to the high consumption of these species and their vulnerability to animal diseases (23). Rwanda has had different zoonotic disease incidences, of which some have become endemic and pose a serious threat to animal and public health. These include brucellosis, Rift Valley fever, rabies, tuberculosis, and cysticercosis (24-27). A study was done in Rwanda on women with abortion and stillbirth complications in two hospitals of the Huye district that revealed 25.0% brucellosis seropositivity (28). The bovine seroprevalence at the interface of livestock, wildlife, and humans was 7.4% and 28.9% at individual animal and herd level, respectively (13). The studies conducted in Nyagatare district reported that the seroprevalence of bovine brucellosis varies between 1.7 to 18.9% (29, 30). These studies confirmed the endemicity of brucellosis in Rwanda and mentioned that human brucellosis from interaction with animals is a public health threat. However, no research was done on the prevalence of *Brucella* infection in small livestock in Rwanda (30). The study aimed to figure out seroprevalence and examine risk factors connected with seropositivity of *Brucella* spp. in goats raised in Nyagatare district in Rwanda. It is anticipated that the findings of this study will aid in the development of future disease control programmes and interventions.

## Material and methods

### Study area

The study was conducted in Nyagatare district, which is one of the 30 districts in Rwanda (**Fig 1**). The district has 14 sectors organised in 106 cells and 630 villages. Fourteen sectors that made Nyagatare are Nyagatare, Matimba, Rwimiyaga, Rwempasha, Karangazi, Rukomo, Katabagemu, Karama, Kiyombe, Mukama, Tabagwe, Mimuli, Gatunda, Musheri. Nyagatare district is bordering Uganda and Tanzania and Gatsibo district. The district covers an area of 1,919 km^2^ and has an altitude of 1414m. The main sectors of sample collection were Nyagatare, Rwempasha, Rwimiyaga, Matimba, Katabagemu and Musheri (**Fig 1**). Nyagatare has a population number of 465,855 (31). The average human density is 243 inhabitants/km^2^ in Nyagatare district compared to the national average human density of 521 inhabitants/km^2^(31).

**Fig 1:**
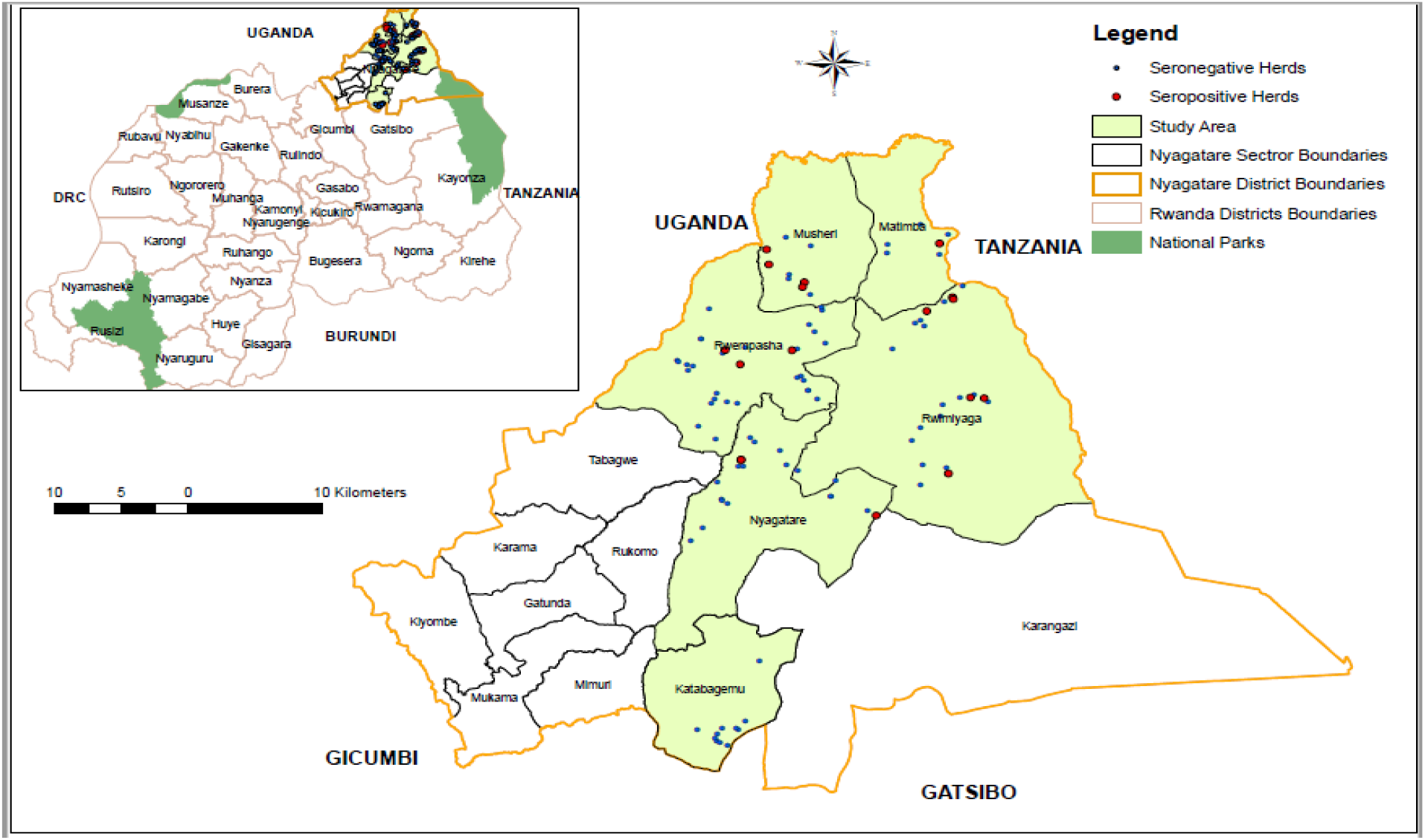
Left, Rwanda map with the location of the Nyagatare district outlined in yellow, Right, Nyagatare district with 14 sectors, and sampled sectors highlighted in green. Red and blue circles show *Brucella* seropositive and seronegative caprine herds found in this study.

### Study animals

Female and male goats older than six months with no brucellosis vaccination history were selected. The district veterinary office report from 2017 to 2018 indicated that Nyagatare district has 63,808 goats (according to district veterinary services records). The goat rearing systems in Nyagatare include free ranching, semi-intensive, and tethering.

### Study design and sampling strategy

A cross sectional study took place from November 2020 to September 2021. Of the 14 sectors of the Nyagatare district, 6 sectors with high goat numbers namely, Nyagatare, Rwempasha, Rwimiyaga, Matimba, Katabagemu, and Musheli sectors were selected. A multistage sampling method was used to sample the herd as the primary sampling unit using the formula for simple random sampling suggested by Thrusfield (32).

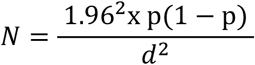

The sample size of the herd was determined using a 95% confidence interval level with an expected prevalence (P) of 13 % of caprine brucellosis at the herd level and with a desired absolute precision (d) of 10%. The formula yielded 44 herds, and the sample size was inflated two-fold to be 88 for the purpose of increasing the precision (33-35). Besides, a contingency of 15% of herds was added, leading to a total of 102 herds sampled in selected sectors of the Nyagatare district. The prevalence of 13% used in sample size determination was obtained from a similar study conducted in Uganda in an area that has the same ecosystem as Nyagatare district (12). The number of herds that were randomly sampled within the sector was based on the proportionate sampling scheme since the total numbers of flocks per sector and district are available from the Nyagatare district veterinary office and formed part of the initial stage sampling. The herd sample size determined was chosen randomly using an Excel random generator from the respective list of herds that belong to different sectors.

Within the herd, all goats above six months of age present during the time were grouped in an enclosed pen, counted, and identified using ear tags. A list of all goats from 1 to N (N= total number of goats eligible to be sampled in the herd) in each herd was made by randomly recording ear tags. Each ear tag was linked to a number from 1 to N according to the first recorded till the last N recorded goat. The next stage of sampling included a fixed number of six recorded goats eligible for criteria of selection that were sampled in each herd by randomly picking numbers in a hat. The total sample size in all 102 herds was 612 goats. From the list obtained in the district veterinary office, 52% of farms are smaller than 5 hectares and goats are mixed with cattle on many farms which limits the number of goats per farm/herd. As suggested by Dohoo (35) and due to financial reasons, to ensure that all animals have the approximate same probability of being selected, a herd below 20 and above 25 goats were excluded from the study and replaced by the following herd on the Excel generated random list which met the herd size criteria.

In this study, a herd was considered as all animals reared in the same household visited. Information on seroprevalence was obtained from laboratory analysis of blood serum. Information on risk factors such as parity, flock size, and abortion was obtained from the head of the selected herd/household using a questionnaire, well arranged, tested, and piloted on 10 goat farms.

### Blood sample collection

The collection of blood was done on the jugular vein of goats, using non-anticoagulant tubes. Blood samples (4ml) collected were transported on the cold chain at 4°C and were kept in the laboratory at room temperature overnight for clotting. The serum was collected in eppendorf tubes, labelled, and stored in the freezer at -20°C until the time of analysis. Laboratory analyses were conducted in the laboratory of University of Rwanda, School of Veterinary Medicine.

### Blood sample analysis

The samples were screened using the Rose Bengal test (RBT), and indirect enzyme-linked immunosorbent assays (iELISA). Both test reagents were obtained from IDvet Diagnostics (France). Overall herd prevalence was determined based on seropositive samples identified on both RBT and iELISA.

#### Rose Bengal test (RBT)

Serum samples were analysed with RBT using an antigen reagent from ID Vet (France). The serum was mixed in equal amounts with the RBT antigen using a plastic plate. Shortly following the final antigen drop, the serum was mixed thoroughly with antigen to produce a round surface that has a diameter of 2 cm. The serum and antigen were mixed in agitating motion for 4 min at the temperature of the room. Visible agglutination reactions were recorded as positive immediately after 4 min of agitation. The sensitivity of the test condition was tested using a positive serum with a minimum positive reaction.

#### Indirect ELISA

All serum samples were analysed using an indirect ELISA kit for brucellosis detection (ID Vet, France) by following appropriately the protocol as designed by the manufacturer. The iELISA uses *Brucella* lipopolysaccharide (LPS) as an antigen to detect anti-*Brucella* antibodies in small ruminant serum. A negative and positive control were included in every test. Samples and the controls were diluted at 1/20 and formed an antibody-antigen complex if present. The horseradish peroxidase (HRP) conjugate added, forms an antigen-antibody conjugate-HRP complex. After washing to eliminate the excess conjugate, the substrate (TMB; 3, 3’, 5, 5’-tetramethylbenzidine) was added, which resulted in a colour reaction depending on the quantity of specific antibodies and measured at 450 nm using a spectrophotometer Multiskan reader (Thermo Scientific, USA). A positive result was obtained when a sample showed S/P greater than or equal to 120 %. S/P%= [(OD sample-OD_Nc_)/(OD_Pc_-OD_Nc_)]x100 with OD = Optic density, Pc = positive control, and Nc= negative control.

### Assessment of risk factors associated with brucellosis

#### Questionnaire administration

A pre-tested questionnaire was utilized to collect information related to risk factors associated with brucellosis. Factors that were examined include mixing small and large ruminants, contact with other herds, management and breeding practices, and the addition of a new goat in the flock in the past year. (**S1 Questionnaire**). The questionnaire in the local language was used to collect information from farmers that participated in the study. The questionnaire had been pretested on ten animal owners to check for consistency and any ambiguity.

#### Risk factor analysis

R Studio and Ms Excel 2013 were used for entering, cleaning, and analyzing data. The univariate analysis was performed where proportions were calculated for categorical variables and means and medians for continuous variables. The bivariate analysis was carried out to evaluate the association between herd *Brucella* seropositivity and potential risk factors. The confidence interval of 95% and odds ratios were calculated, and factors were considered statistically significant if having a p-value of ≤ 0.05.

The p-value ≤ 0.20 criterium was used to select independent variables for inclusion in the initial multiple logistic regression model. The model was constructed using a stepwise selection procedure, dropping the least significant independent variable and continuing until all remaining predictor variables were significant (p-value ≤ 0.05).

### Ethics approval and protocol

Assessment and approval of the study protocol were done by the University of Rwanda, the University of Pretoria, the Faculty of Veterinary Science and Faculty of Humanities. In each cell, goat farmers held a meeting to discuss the aim of this research. Study participants were not obliged to participate in the study or to allow the collection of the blood of their animals. The questionnaire was elucidated to the participants, and they were encouraged to ask questions for more clarification. Participants that provided verbal consent to partake in the questionnaire and allowed blood sampling of animals were enrolled in the study. The farmers’ names, regions, and villages were recorded in a non-disclosure agreement. The questionnaire used in the survey was coded and saved in Excel. For analysis, codes were utilized to substitute farmer names. Blood samples were collected by qualified veterinarians in accordance with animal husbandry regulations.

## Results

### Sero-prevalence of brucellosis at individual and herd level

#### Herd characteristics

A total of 102 herds distributed in 6 different sectors in Nyagatare district (**Fig 1**) were visited for samples collection and questionnaire administration. In this district, 95 out of 102 respondents (93.1%) kept mixed herds (thus goats with other animals), while 3.9 % of respondents kept their goats and other animals separately. The goat herds were 87.3%, 54.9%, 47.1%, and 30.4% mixed with cattle, dogs, poultry, and sheep, respectively. Of the respondents, 82.9% of the goats shared pastures and watering places with other animals. 612 goats were sampled from 102 herds in the district.

#### Brucellosis seroprevalence at an individual animal level

Of the 612 goat samples, 6.8% (42/612) were RBT brucellosis seropositive and 10.7% (66/612) were iELISA seropositive. All the RBT seropositive samples were positive with iELISA and thus 6.8% were seropositive using iELISA and RBT in Nyagatare district. An additional 24 samples tested iELISA positive but RBT negative (**Table 1**).

**Table 1:**
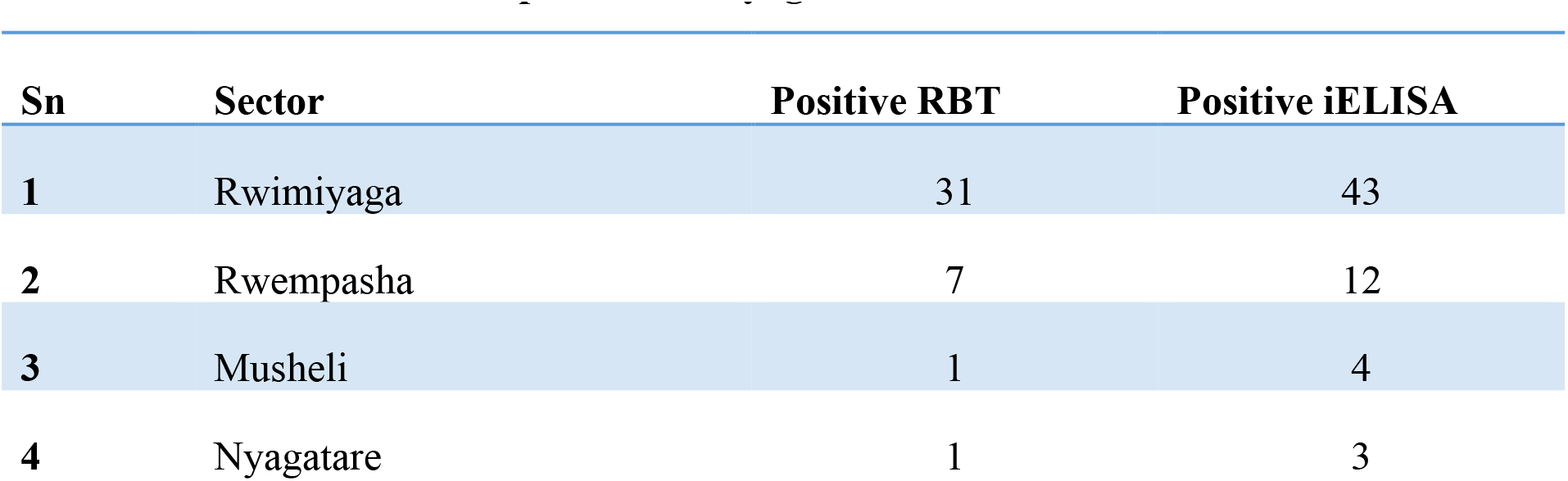
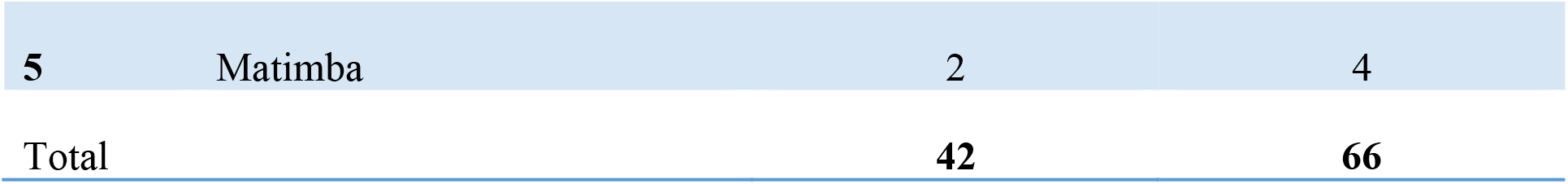
Brucellosis seroprevalence using RBT and iELISA at the individual animal level in different sectors sampled in the Nyagatare district.

#### Brucellosis sero-prevalence at herd level

The herd was confirmed to be positive when at minimum one goat in the herd was confirmed to be positive for both RBT and iELISA with 16.6% (17/102) herds positive in the Nyagatare district **(Table 2)**.

**Table 2:** Brucellosis positive animals in herds in Nyagatare district using RBT and iELISA.

| The number of goats seropositive in the flock | Number of flocks with positive reactors on both RBT and iELISA | Number of flocks with positive reactors using iELISA |
| --- | --- | --- |
| 1 | 6 | 8 |
| 2 | 4 | 5 |
| 3 | 1 | 2 |
| 4 | 4 | 5 |
| 5 | 2 | 3 |
| 6 | 0 | 1 |
| Total | 17 | 24 |

### Factors associated with brucellosis herd seropositivity

#### Bivariate analysis for herd level factors linked to brucellosis in goats

In determining herd level factors linked to brucellosis in goat herds, different factors were analysed to detect their association with seropositive herds **(Table 3)**. Only three risk factors among 11 were statistically associated with herd seropositivity. The majority of study respondents (66.6%) mix their cattle and goat herds and that was recognized as among the risk factors this study had identified. Mixed goat and cow herds had an eightfold increased risk of having positive brucellosis reactors when related to goat herds that were not mixed with cattle herds and thus statistically significant (OR=8; p=0.049) **(Table 3)**.

**Table 3:**
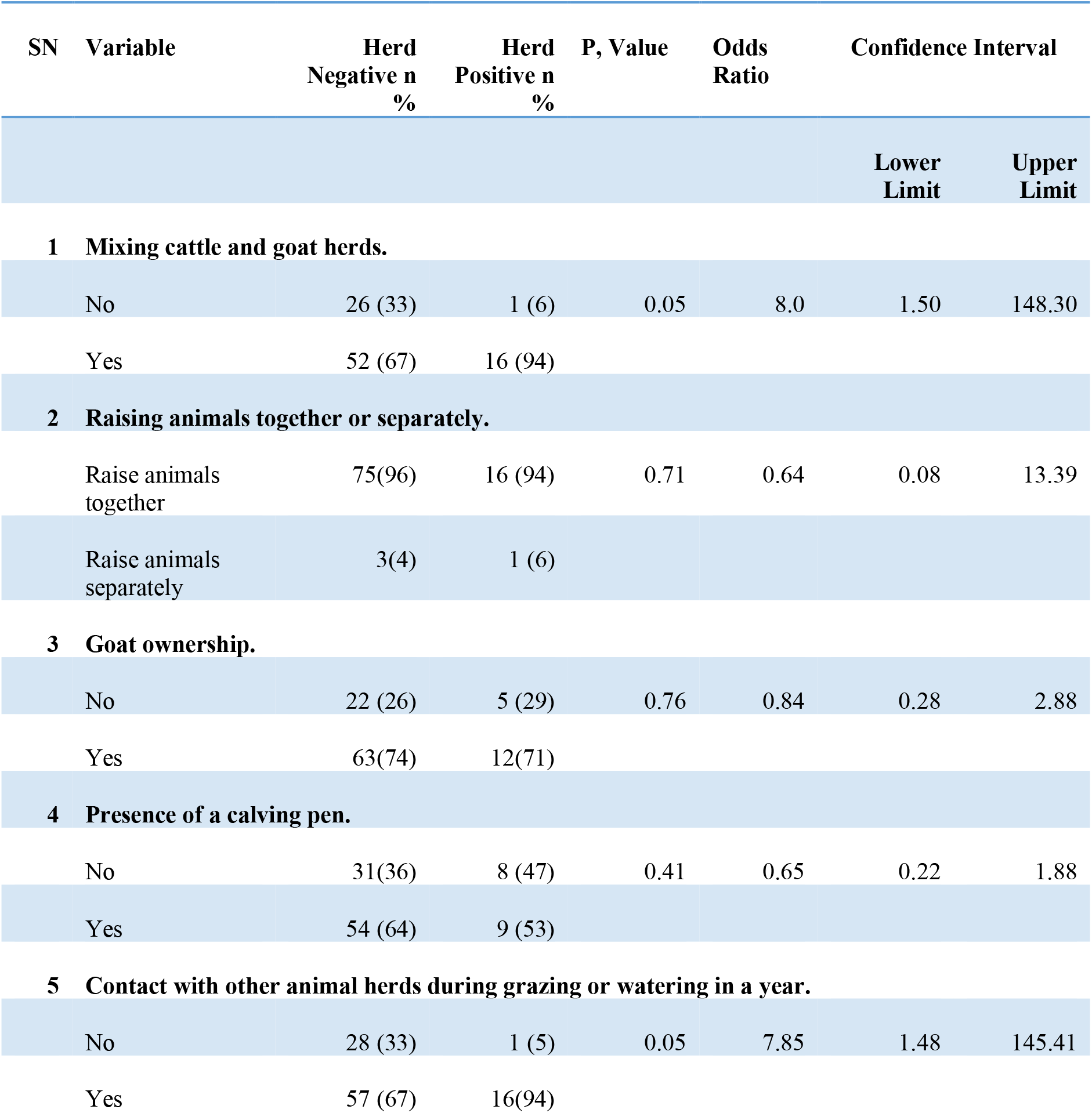
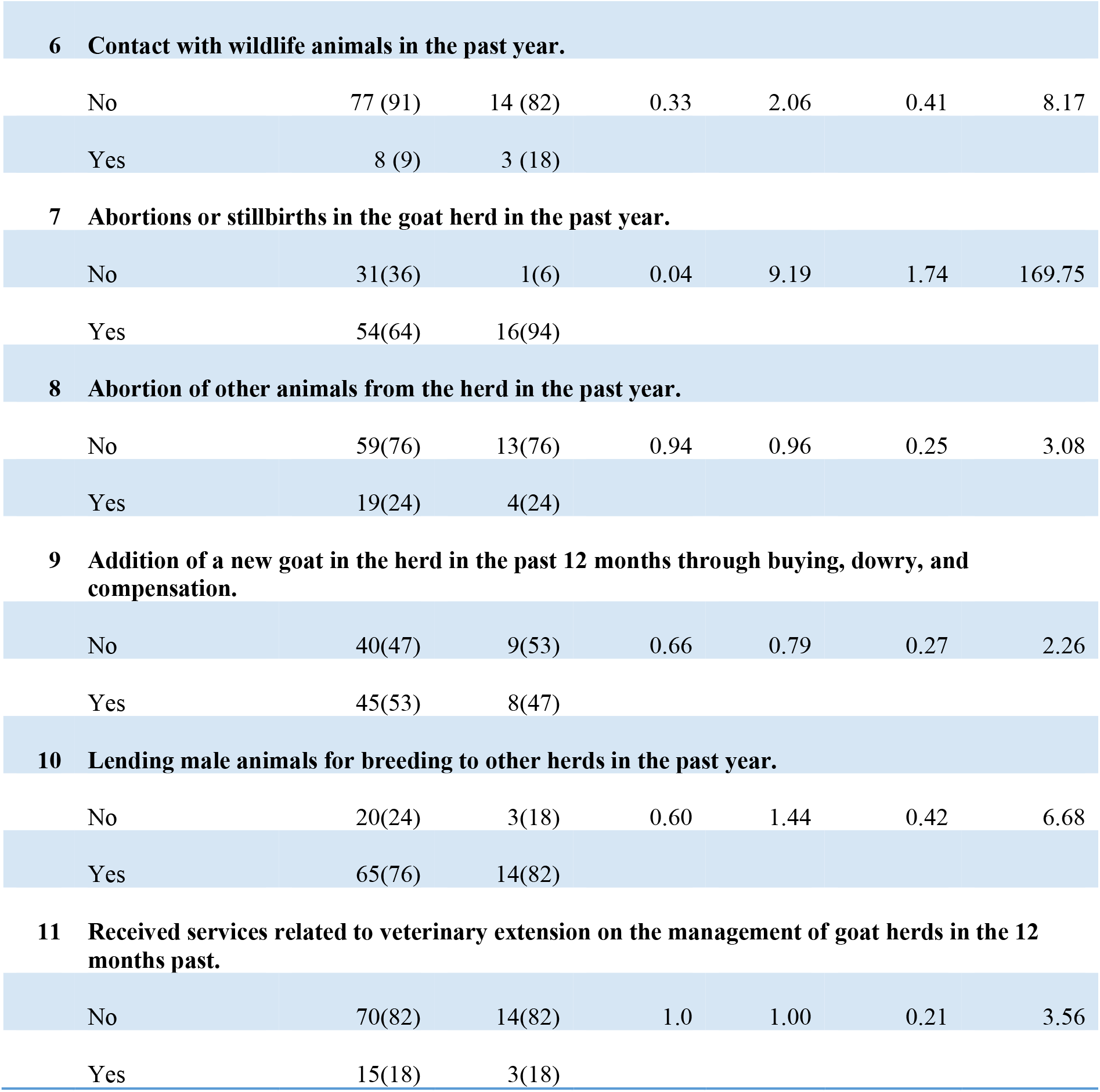
Comparison of risk factors associated with brucellosis seropositive and seronegative herds in Nyagatare district.

Abortions were reported in 70/102 (68.6%) goat herds. Of the seropositive herds, 16/17 (94.1%) had experienced abortion compared to 64.0% (54/85) abortions reported in seronegative herds. Herds with an abortion history were nine-fold increased chance of having seropositive animals when put in comparison to herds with no abortion history, and thus statistically significant (OR = 9; p = 0.035). However, the history of abortion in other animal herds (OR= 0.95; p-value = 0.940) was not linked with *Brucella* positive goat herds **(Table 3)**.

Interaction with other goat flocks when grazing or watering was indicated in 71.5% of the visited herds. Goat herds that interacted with other herds during grazing or watering were eightfold more chances to be positive for brucellosis than herds without contact with other goat herds and was statistically significant (OR=7,85; p=0.05). Contact of goat herds with wildlife animals was not statistically associated with brucellosis seropositive herd (OR= 2; p= 0.320) **(Table 3)**.

All respondents did not vaccinate their animals and the majority of study participants (82.4%) did not get veterinary advice on the management of their goats. A small proportion of the respondents (17.7%) indicated that they quarantined new animals before introducing them into the herd.

#### Multivariable analysis to determine independent factors with *Brucella* herd seropositivity

The multivariable logistic regression model revealed that mixing cattle and goat herds (adjusted odds ratio (aOR) = 8.35; p=0.04) and abortion history in the goat herds (aOR =0 7.94; 95%; p=0.05) were the independent factors associated with goat herds seropositivity to *Brucella* spp. The odd value of contact between goat herds during grazing and watering was statistically insignificant ((aOR) = 6.00; p-value 0.09) with herd seropositivity **(Table 4)**.

**Table 4:**
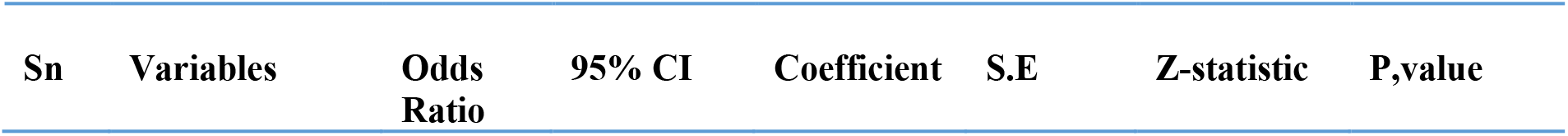
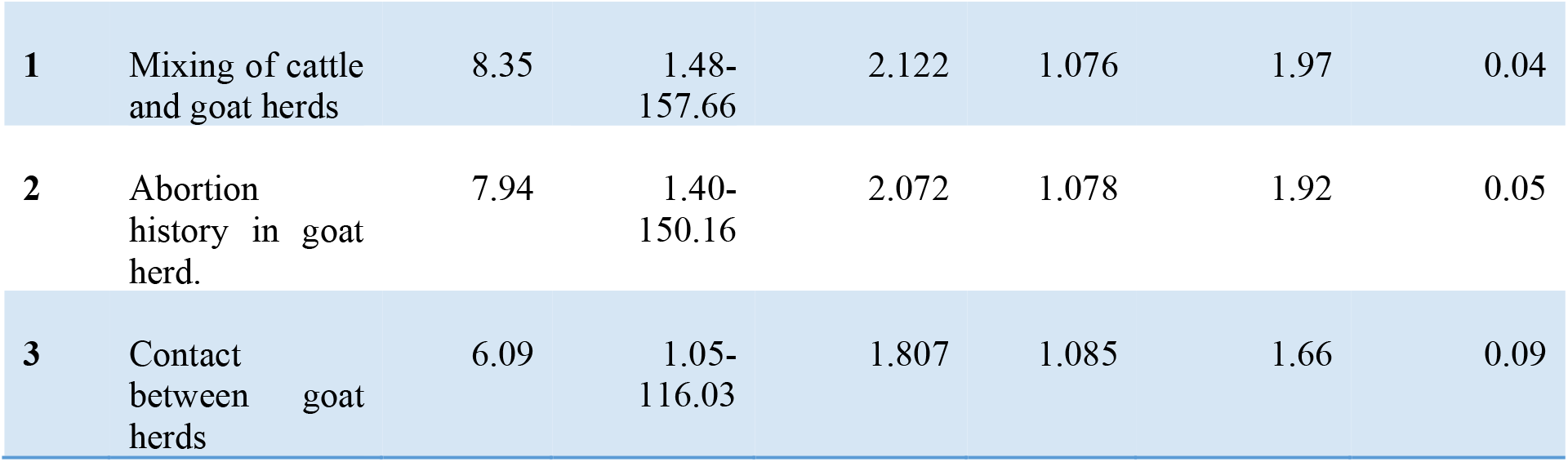
Analysis of the variables associated with herd-level seropositivity for brucellosis in goat herds in Nyagatare district using multivariable logistic regression.

## Discussion

### Sero-prevalence of brucellosis in goats

The brucellosis seroprevalence at individual goat level was 6.8% and 10.7% on RBT and iELISA consecutively. All seropositive samples on RBT were confirmed to be seropositive to iELISA. The difference in the number of seropositive samples between the used tests was also recorded in the findings of Kishan in India who reported 49 serum samples that were positive in iELISA but negative in RBT (36). High number of seropositive animals recorded with iELISA that was seronegative for RBT might be attributed to the type of antibody present in the serum (IgM, IgG_1_, and IgG_2)._ The iELISA ability to detect antibody response against brucellosis persists after the peak 3 to 4 weeks postinfection and can remain diagnosed for many years, while IgM response for brucellosis mainly detected by RBT is rapidly induced and may disappear after a short period (37-43).

The cut-off value of iELISA needs to be validated in Rwanda as the use of current cut-off point may underestimate brucellosis in caprine in Rwanda as it is based on kits developed in countries with low brucellosis prevalence. Furthermore, it is important to determine if iELISA positive and RBT seronegative animals are chronic infections of brucellosis. This can be confirmed by diagnostics methods such as culture. The discrepancy in the number of *Brucella* positive cases discovered in this study using the RBT and iELISA tests can also be linked to the specificity and sensitivity of each test (44, 45). *Yersinia enterocolitica O: 9* as a cross-reactive bacterium can also contribute to the difference in the number of seropositive animals identified by RBT and iELISA (16). All positive results obtained in this study are not due to vaccination as all respondents did not vaccinate their goats. Furthermore, there is not any government campaign for brucellosis vaccination in Rwanda in small ruminants.

Studies conducted in Rwanda indicated that the seroprevalence of bovine brucellosis in the last decade varied between 1.7% and 18.9% among cattle in Rwanda. (13, 29, 30, 46). The high record of seroprevalence of bovine brucellosis of 18.9% is in the Nyagatare district (29). The seroprevalence of bovine brucellosis in the Nyagatare district is higher compared to the recorded seroprevalence of caprine brucellosis in the same district. This may be attributed to the sampling strategy used and the poor capacity of the dairy farms to keep bulls, which lead to a high exchange of bulls between farms for mating thus the easy spread of brucellosis among cattle.

The study reported brucellosis seroprevalence in goats of Nyagatare district, which differ from the results of caprine brucellosis seroprevalence in Ethiopia. Those studies screened sera goat samples using RBT and did confirmation with CFT or iELISA and indicated the seroprevalence at individual animal and herd level, varying between 0.4% to 9.7%, and 15% to 45% consecutively (47-51). In Niger, brucellosis seroprevalence was reported to be between 0.4% and 3.6% individual animal level and 17.8% herd level using iELISA (52). A reviewed study on seroprevalence of *Brucella* infection in goats in the neighbouring country of Rwanda (Burundi, Kenya, Uganda, Tanzania, and South Sudan) reported varying seroprevalence of 0.0% to 20.0%. The studies done in Uganda revealed caprine brucellosis seroprevalence between 0.3%, and 14.82% using iELISA (53, 54) Another study in Kiruhura district of Uganda revealed 8.8% seroprevalence in goats using RBT (55). Studies done in Kenya using cELISA estimated individual animal-level seroprevalence ranging from 1.3% to 16.1% and herd-level seroprevalence ranging from 5.6% to 6,8% (56). Brucellosis seroprevalence of 7.3% and 13.04% in goats were also reported in Kenya using RBT and cELISA as screening and confirmatory test (57, 58). In Tanzania, brucellosis seroprevalence of 10.7% bucks, and 11.9 % of does were reported by a study done in livestock-wildlife-interface at Ngorongoro conservation area using RBT (59). Furthermore, the comparison between the seroprevalence of caprine brucellosis in Rwanda and neighbouring countries indicates variation, which can be attributed to different aspects including systems of production, animal husbandry, and a variety of agroecological zones in which the studies were carried out. It can also be attributed to the sampling methodology and laboratory test used in the identification of seropositive serum. The mentioned factors were also characterized by other researchers to contribute to the variation of brucellosis seroprevalence. (39, 60-62).

### Factors at the herd level that influence brucellosis in goats

Abortion records in the goat flock were found to be significantly linked to *Brucella* seropositivity in the herd. Abortion in small ruminants is the major clinical sign of brucellosis infection (12, 63, 64). Female animals infected by brucellosis release a high concentration of *Brucella* spp. in their milk, placenta, and aborted materials (9). The release of *Brucella* microorganisms by infected female animals occurs for a long period, which results in the contamination of the environment (5, 65). The risk factors for brucellosis may be increased by respondents’ attitude to discard the aborted fetus in the environment, which results in an increased probability of transmission of brucellosis within or between herds during grazing and in watering places. Furthermore, the history of abortion characterized by this study as the main risk factor, can favour zoonotic transmission of brucellosis, as the majority of respondents reported to assist their animals during parturition or handled aborted materials without any protective clothing. Respondents also declared to drink raw milk, and those attitudes put them at high risk of being contaminated in case those materials are contaminated. The study indicated a high rate of abortions in herds, 68.3% this findings agree with what reported by Mourad (66) on the high abortion rate of goats in Rwanda. The history of abortions recorded in 64% of seronegative herds can be attributed to different unidentified causes which may include endemic zoonotic diseases like Rift Valley Fever (26).

Raising cattle herds and goat herds together was characterized as significantly connected with herd seropositivity. This finding agrees with the research done in Ethiopia, Egypt, and Rwanda which confirmed mixing cattle and goat herds as a risk factor linked to seropositive herds of cattle (13, 47, 67). Furthermore, the findings in this study are in agreement with study findings published on the risk factors for bovine brucellosis in Uganda and Tanzania (68, 69). A study conducted in Kenya also revealed the transmission of *Brucella abortus* between cattle and goats (70). Transfer of brucellosis infection between cattle and goat herds could be easily facilitated by the improper disposal of an aborted fetus (65, 71) and the fact that there is no vaccination campaign in Rwanda against brucellosis in both large animals and small animals (72).

Contact with other herds during grazing or watering was characterized to be statistically associated with herd seropositivity in bivariate analysis, which was also reported in Niger and Kenya (11, 52). However, it is in contrast with studies done in Uganda and Tajikistan (12, 73). The findings on contact between goat herds obtained in Rwanda might be considered as a result of a poor understanding of the transmission mode of brucellosis in animals as well as the lack of a brucellosis control program. Other contributing factors might be the lack of fences on some of the farms, which allow free movement of goats and other animals between farms (74). Besides, the shortage of water in the Nyagatare district causes the herds of goats to share a communal watering point (74, 75).

The study revealed that lending breeding males and new animals introduced into the herd without quarantine were not significantly associated with herd seropositivity which contrasts with similar studies conducted in different places, including neighbouring countries Uganda, and Kenya (11, 12). This can be attributed to the fact that farmers have the attitude of lending males from a neighbour where they know well that there is no history of abortions in the herd. Veterinary advice on the management of goats and vaccinating animals against brucellosis were assessed as preventive factors during this study and were not significantly associated with herd seropositivity. Contrasting results were recorded in the study conducted in Uganda and Kenya where veterinary service is considered as linked with herd seronegativity as a protective factor (11, 12). This may be attributable to the reality that nearly every study participant didn’t get either veterinary advice in the management of goat herds or vaccinate their goats.

Only three risk factors were associated with caprine brucellosis in this study, while the following ten risk factors were characterized to be associated with bovine brucellosis in Rwanda. The use of an open grazing system, a history of abortion, a history of longer calving intervals (> 1 year), and the retention of the placenta were all significant risk factors for the presence of anti-Brucella antibodies in cow milk(76). Old age (> 5 years), cattle farmed close to wildlife, herds of cattle mixed with small ruminants, history of abortions, and replacement animals were significantly associated with brucellosis(13). Age, number, breed, and parity were also identified as risk factors for bovine brucellosis in the Nyagatare district(29). The difference in risk factor associated with brucellosis between goat and cows may be attributed to the fact that few study was conducted on caprine brucellosis. A history of abortion and mixed herds of cattle and goats on the same farm were described as risk factors for brucellosis in both goats and cattle herds in Rwanda, and this may be attributed to the fact that animals with a history of abortion are not removed on the farm and the fact that in the Nyagatare district, the majority of farms have cattle and goats mixed and raised in the same conditions where they share pasture, shelter, and water.

## Conclusions and Recommendations

The study confirmed the endemicity of brucellosis in the Nyagatare district, which was demonstrated by the outrageous seroprevalence of brucellosis at both individual animal and herd level. There is a high potential for transmission of brucellosis within the ecosystem to all susceptible candidates. Mixing herds of cattle and goats, as well as the history of abortion, were characterised to be the major independent risk factors associated with herd seropositivity. In this regard, the study suspects easy transmission of brucellosis within mixed herds or between herds. The majority of the community is not aware of the zoonotic aspect of brucellosis and doesn’t know that brucellosis affects goats. They are also highly involved in practices that can lead to the spread of brucellosis within-herd and between herds without living behind humans. The study recommends establishing a national contingency programme for the control and prevention of brucellosis. It is paramount to raise awareness of brucellosis and other zoonotic diseases in the community and help establish prevention and control measures. One health approach is highly recommended to control and prevent brucellosis in the Nyagatare district. The study recommends an awareness campaign for biosafety measures when handling aborted fetuses and further studies are recommended to have all necessary data on brucellosis in the ecosystem.

## Acknowledgements

The authors would like to acknowledge the Institute of Tropical Medicine, Antwerp, Belgium (ITM)), the University of Pretoria (UP) for all support to conduct this study, and the University of Rwanda, School of Veterinary Medicine for the facilitation of this study. We also acknowledge the District Veterinary Officer and Sector Veterinary Officers in Nyagatare district for helping me with access to farms and helping me in organizing the sampling process.

## Author contribution

**Conceptualisation**: Jean Paul Habimana,

**Data curation** : Jean Paul Habimana,

**Laboratory analysis**: Jean Paul Habimana,

**Methodology**: Jean Paul Habimana and Prof. Henriette van Heerden,

**Investigation:** Jean Paul Habimana, Eric Gasana, and Aurore Ugirabe

**Statistical analysis**: Jean Paul Habimana, Aime Lambert Uwimana, and Eric Gasana

**Supervisor:** Prof. Henriette van Heerden.

**Writing -original draft:** Jean Paul Habimana

**Writing-review and editing:** Jean Paul Habimana, Dr Jean Bosco Ntivuguruzwa, And Prof. Henriette van Heerden,

## Supporting information

S1 Questionnaire. Risk factor Study—Brucellosis in Nyagatare district.(DOC)

S1 Data. Raw data in excel format.(XLSX)

